# Response of coral reef dinoflagellates to nanoplastics under experimental conditions

**DOI:** 10.1101/2020.09.23.310847

**Authors:** Christina Ripken, Konstantin Khalturin, Eiichi Shoguchi

## Abstract

Plastic products contribute heavily to anthropogenic pollution of the oceans. Small plastic particles in the micro- and nanoscale ranges have been found in all marine ecosystems, but little is known about their effects upon marine organisms. In this study we examine changes in cell growth, aggregation, and gene expression of two symbiotic dinoflagellates of the family Symbiodiniaceae, *Symbiodinium tridacnidorum* (clade A3) and *Cladocopium* sp. (clade C), under exposure to 42-nm polystyrene beads. In laboratory experiments, cell number and aggregation were reduced after 10 days of nanoplastic exposure at 0.01, 0.1, and 10 mg/L concentrations, but no clear correlation with plastic concentration was observed. Genes involved in dynein motor function were upregulated compared to control conditions, while genes related to photosynthesis, mitosis, and intracellular degradation were downregulated. Overall, nanoplastic exposure led to more genes being downregulated than upregulated and the number of genes with altered expression was larger in *Cladocopium* sp. than in *S. tridacnidorum*, suggesting different sensitivity to nanoplastic between species. Our data show that nanoplastic inhibits growth and alters aggregation properties of microalgae, which may negatively affect the uptake of these indispensable symbionts by coral reef organisms.

## 1. Introduction

Coral reefs provide habitat for marine invertebrate and vertebrate species alike, sustaining the highest biodiversity among marine ecosystems [1]. Formed primarily by scleractinian corals and coralline algae, coral reefs are complex and vulnerable ecosystems. Structural complexity of coral reefs, and by extension, the capability to sustain biodiversity often declines due to natural and human-related stressors [2,3].

One important stressor for coral reef ecosystems is plastic pollution. Small plastic particles (>1 mm) have been reported from coral islands at more than 1000 items/m^2^ [4]. Further fragmentation of these particles leads to nanoplastics (<1 μm) [5]. Microplastic particles induce stress responses in scleractinian corals, suppress their immune systems and capacity to cope with environmental toxins [6]. When ingested by corals [7,8,9], microplastics disrupt the anthozoan-algal symbiotic relationship [10]. They are also linked to potential adverse effects on calcification [11] with exposure resulting in attachment of microplastic particles to tentacles or mesenterial filaments, ingestion of microplastic particles, and increased mucus production [12]. Su *et al*. [13] exposed the coral symbiont, *Cladocopium goreaui*, to 1-μm polystyrene spheres, leading to diminished detoxification activity, nutrient uptake, and photosynthesis, as well as increased oxidative stress, apoptosis levels, and ion transport. Plastic particles seem to negatively impact symbiotic relationships between corals and their microalgae, thereby degrading the entire coral reef ecosystem, but this has not been systematically investigated.

Nanoplastics, particles smaller than 1 μm [5], can originate by fragmentation of larger plastic objects through photochemical and mechanical degradation. There are also primary sources of nanoplastics. Medical and cosmetic products, nanofibers from clothes and carpets, 3D printing, and Styrofoam byproducts find their way into coral reef ecosystems via river drainages, sewage outfalls, and runoff after heavy rainfall, as well as via atmospheric input and ocean currents. Nanoplastic has recently been reported in ocean surface water samples [14].

In this study we focused on the microalgal symbionts of mollusks that inhabit fringing coral reefs of Okinawa. Knowledge of the effects of nanoplastic on the symbionts of Tridacninae (giant clams) and Fraginae (heart cockles) will benefit conservation and restocking efforts, as both are obligatory photo-symbionts and important contributors to coral reef ecosystems. Approximately 30 Symbiodiniaceae phylotypes are economically important for fisheries [15]. This study specifically investigated effects of nanoplastic (42-nm polystyrene spheres) on the growth rates, aggregations, and gene expression changes in *Symbiodinium tridacnidorum* (symbionts of the Tridacninae) and *Cladocopium* sp. (symbionts of the Fraginae).

## 2. Materials and Methods

### 2.1. Exposure to nanoplastics using roller tanks

The majority of host animals obtain their indispensable symbiotic dinoflagellates from coral reef sand and the water column [16, 17]. Roller tanks and tables were used to simulate the natural environment of the dinoflagellate vegetative cells in their free-living state [18, 19]. Roller tanks have commonly been used to promote aggregation since Shanks and Edmondson [19, 20]. 15 roller tanks 13.4 cm in diameter and 7.5 cm in height with a capacity of 1,057 mL were employed. In tanks, aggregation can occur [19], ensuring that microalgae are exposed to the polystyrene nanoplastic (nanoPS) in a way that mimics their natural habitat. Once rotation commenced, continuous aggregate formation and suspension were ensured [20] as well as continuous exposure to nanoPS. Roller tanks are closed for the entire duration of the experiment, so that exposure levels of the nanoPS remain constant through-out. Tanks were closed without bubbles so as not to disturb the aggregation process with turbulence. To compare differences between species, two dinoflagellates, *Symbiodinium tridacnidorum* (clade A3 strain, ID: NIES-4076) and *Cladocopium* sp. (clade C strain, ID: NIES-4077) were cultured in artificial seawater containing 0.2x Guillard’s (F/2) marine-water enrichment solution (Sigma-Aldrich) in roller tanks [21,22]. *S. tridacnidorum* and *Cladocopium* sp. (Clade C strain ID: NIES-4077) were isolated from *Tridacna crocea* and *Fragum* sp. in Okinawa, Japan [5]. Using glass flasks, precultures for the stress experiment were established, as previously described [4].

Microplastics (>1 mm) from coral reef and the ingestion (53 to 500 μm) by coral reef clams have been reported and microplastic removal by giant clams has been proposed [4, 23]. To simulate nanoplastic accumulation in coral reefs and in the host organisms, three different concentrations (0.01 mg/L, 0.1 mg/L, and 10 mg/L) of nanoplastic (42-nm pristine polystyrene beads, nanoPS_42_, from Bangs Laboratories Inc., catalog number FSDG001, polystyrene density 1.05 g/cm^3^, nanoPS) were added to the treatment tanks (Tables S1). Treatment tanks as well as control tanks (no nanoPS) were established in triplicate. Three tanks without algae were prepared as negative controls (at 10 mg/L, 0.01 mg/L, 0 mg/L nanoplastic). In each culture tank, the final cell density of the two strains was adjusted to ~7 x 10^5^ cells/mL. Tanks were harvested after 9-11 days, for logistical reasons, making replicates a day apart (Supplementary Table 2).

### 2.2. Measurements of cell density and aggregation

Cells for growth rates were counted using hemocytometers (C-Chip DHC-N01) under a Zeiss Axio Imager Z1 microscope (Jena, Germany). At least 2 subsamples and 200 cells were counted per sample.

Aggregates were imaged and counted in each tank and for five size classes, as follows: tiny: 0.2 – 0.5 mm; small: 0.5 – 1 mm; medium: 1 – 2.5 mm; large: 2.5 – 3.5 mm; huge: > 3.5 mm in the longest dimension. Tanks of the same concentration were sampled at the same time of day. Controls were sampled first and then in order of increasing nanoPS_42_ concentration to avoid nanoplastic carry over from higher concentrations to lower. In order to examine how nanoPS_42_ affects aggregate formation, aggregates were collected for different measurements, after the approximate total number off aggregates in each tank had been determined. Aggregation of algae and plastic was confirmed with 3D imaging using a Zeiss Lightsheet Z.1 and Imaris software. NanoPS_42_ was observed with a BP filter (excitation: 405 nm; emission: 505-545 nm) and chloroplasts were visualized using a long-pass red filter (excitation: 488 nm, emission: 660 nm).

One fourth of all aggregates were collected for RNA analysis (2 min spin down at 12,000 rpm and discarding the supernatant, freezing in liquid nitrogen and storage at −80°C). For all other measured factors, harvest included separate sampling of the aggregate fraction (aggregates >0.5mm, Agg) and the surrounding sea water fraction (aggregates <0.5 mm and un-aggregated cells, SSW) [24]. Aggregates for sinking velocity (three aggregates per size class for 11.5 cm in a 100-mL glass graduated glassware cylinder) was collected in artificial seawater at the same temperature as experiments were conducted.

### 2.3. RNA extraction, library construction, and sequencing

Frozen cells were broken mechanically using a polytron (KINEMATICA Inc.) in tubes chilled with liquid nitrogen. RNAs were extracted using Trizol reagent (Invitrogen) according to the manufacturer’s protocol. The quantity and quality of total RNA were checked using a Qubit fluorometer (ThermoFisher) and a TapeStation (Agilent) respectively. Libraries for RNA-seq were constructed using the NEBNext Ultra II Directional RNA Library Prep Kit for Illumina (#E7760, NEB). Sequencing was performed on a NovaSeq6000 SP platform. Nine mRNA-seq libraries from nanoPS-exposed photosymbiotic algae were sequenced (3 concentrations x 3 exposure times) plus three controls (Supplementary Table S2).

### 2.4. RNA-seq data mapping and clustering analysis

Raw sequencing data obtained from the NovaSeq6000 were quality trimmed with Trimmomatic (v0.32) in order to remove adapter sequences and low-quality reads. Paired reads that survived the trimming step (on average 92%) were mapped against reference transcripts of *Symbiodinium* and *Cladocopium* sp.. For each gene in the genomes of *Symbiodinium* and *Cladocopium sp*. a *.t1 transcript form was used as a reference sequence. Mapping was performed using RSEM [25] with bowtie (v1.1.2) as an alignment tool. Expression values across all samples were normalized by the TMM method [26]. Genes with differential expression (2-fold difference and p<0.001) were identified with edgeR Bioconductor, based on the matrix of TMM normalized TPM values. Experimental samples were clustered according to their gene expression characteristics using edgeR. Annotations were performed using BLAST2GO and Pfam databases [21] and are available at the genome browser site (https://marinegenomics.oist.jp).

## 3. Results and Discussion

### 3.1. Suppression of algal growth by nanoplastic exposure

Exposure to nanoPS_42_ decreases the mean growth rate of photosymbiotic algae (see Figure 1). The greatest reduction in growth rate was seen at the lowest nanoPS_42_ treatment (0.01 mg/L), with cell densities reduced from starting values by −0.062 ± 0.02 (Holm-Sidak, p = 0.002); followed by the highest nanoPS_42_ treatment (10 mg/L) with −0.013 ± 0.05 (Holm-Sidak, p = 0.026). In the 0.1 mg/L treatment, cell densities increased slightly by 0.028 ± 0.04. Thus, nanoPS_42_ either inhibited algal growth in a non-linear manner or had a limited effect [27]. Reductions in growth rates have also been reported in the μP study of [13] in *Cladocopium goreaui* and in other microalgae exposed to μP (*Chlamydomonas reinhardtii* [28] and *Skeletonema costatum* [29]).

**Figure 1.**
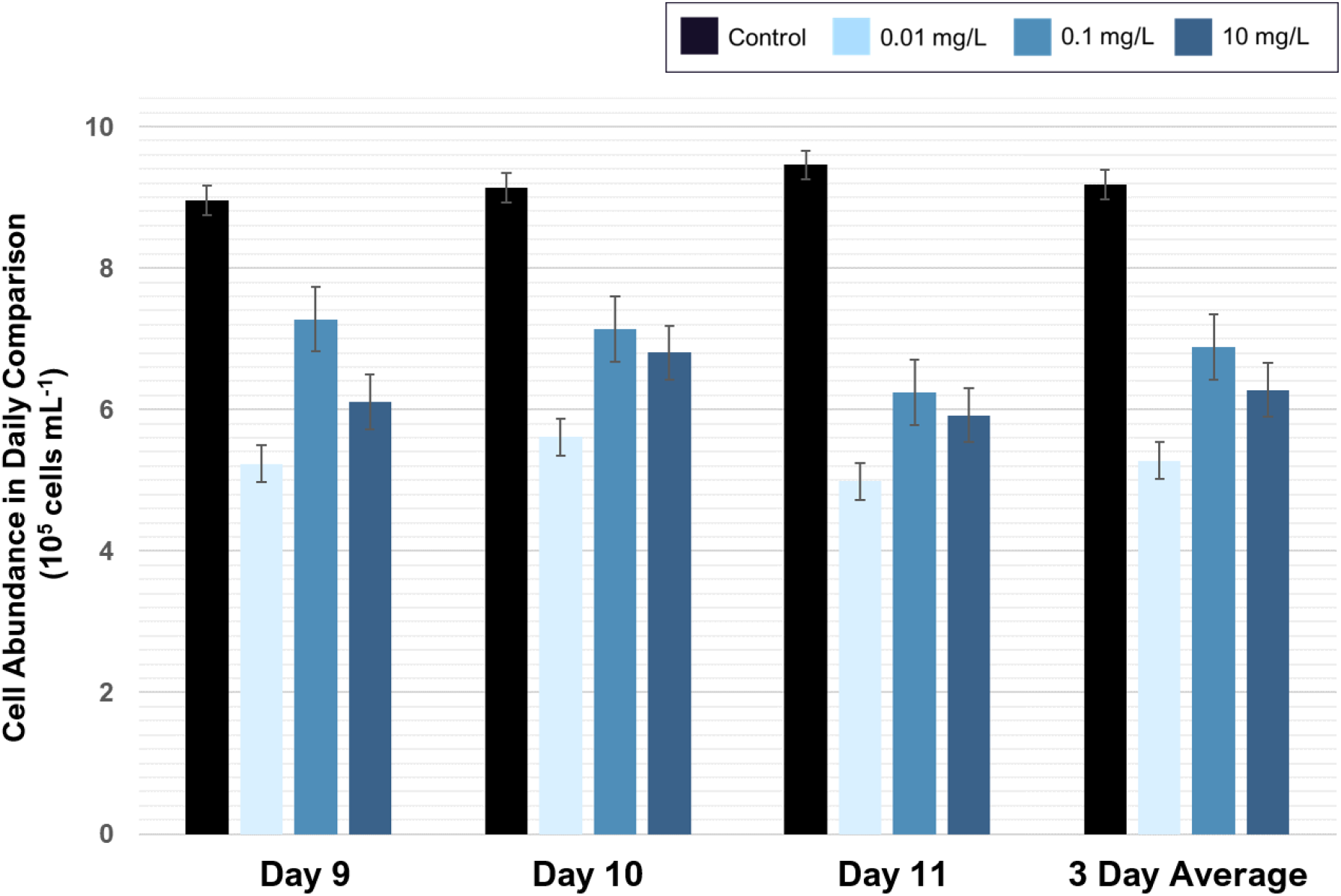
Treatment and control tanks were sampled after 9, 10, and 11 days. Experiments started with ~680,000 cells/mL in all tanks. There are differences between the growth rate in the different treatments, but the ratio stays the same over all three sampling days. The cell density in the control was 9.83 ± 0.39 × 105 cells per mL, while treatment tanks were significantly lower: 0.01 mg/mL: 5.69 ± 0.12 × 105 cells per mL; 0.1 mg/mL: 7.51 ± 0.34 × 105 cells per mL; 10 mg/mL: 6.96 ± 0.40 × 105 cells per mL. Bars display confidence interval.

In addition, Su et al. [13] reported a reduction in cell size in *Cladocopium goreaui*. Further investigations are needed to see if this is the case under nP exposure. Interesting to note is that the biggest growth rate reduction observed was at 0.01 mg/L nanoPS_42_, far below the 5 mg/L used by Su et al. [13]. The nutrient deficiency is also a reason discussed in (Long2017) which could explain the larger effects on growth rates at lower concentrations. The reason for nutrient limitation induced by plastic is proposed to be interactions of the nutrients with the surface of the plastics [30]. NanoPS_42_ self-aggregation could account for the higher nanoPS_42_ treatments having less effect on the growth rates.

### 3.2. Nanoplastic exposure influences the number and sinking velocity of cell aggregates

To understand the impact of nanoPS_42_ on aggregation in these two Symbiodiniaceae cultures, the total number of algal aggregates per tank and in five aggregate size classes was recorded (Supplementary Figure S2). All tanks showed aggregation, which was expected, as self-aggregation of Symbiodiniaceae has been observed previously [13].

The majority of aggregates exhibited an ovoid form. Significant difference can be observed when aggregate numbers are compared over all size classes and all treatments, showing that the nanoPS has an influence on the aggregation process. The lowest nanoPS treatments (0.01 mg/L) shows significant reduction in the total aggregates count by 10 % (Holm-Sidak, p = 0.003). While there is also a reduction of 3 % in the intermediate nanoPS treatment (0.1 mg/L), this is not significant. The different aggregate sizes classes show significantly different distributions in all three treatments and the control (ANOVA, p < 0.001) (Supplementary Figure S2). In the control, the self-aggregation led to a specific distribution pattern of aggregate sizes, which was not repeated in the treatments. Self-aggregation was also observed in the μP experiments of Su et al. [13]. The fact that presence of nanoPS changes the aggregation between the cells and leads to more aggregates in the bigger size classes is possible due to higher production of extracellular polymeric substances (EPS) with sticky properties, trapping more cells in one aggregate and keeping aggregates closer together. Nutrient depletion, which has been linked to the presence of μP in algae cultures [30], is associated with increased stickiness of the extracellular polymeric substances (EPS) [31,32]. Differences in the EPS production due to the presence of nanoPS is a likely factor contributing to the differences in aggregation seen in the study. EPS production was not measured, so further studies are needed to confirm this hypothesis linking the aggregation process and EPS production in Symbiodiniaceae under nanoPS influence. Lagarde et al. [28] notices different aggregate formation under different plastic treatment and sizes, which matches with our results.

Significant differences are evident when aggregate numbers are compared over size classes and treatments, showing that nanoPS influences aggregation. Aggregate size classes show significantly different distributions in all three treatments vs. controls (ANOVA, p < 0.001) (see Figure 2). These differences in aggregation could be due to changes of the cell surface receptors, as nanoPS increases genes related to those 2 fold (see Section NanoPS effects on gene expression).

**Figure 2.**
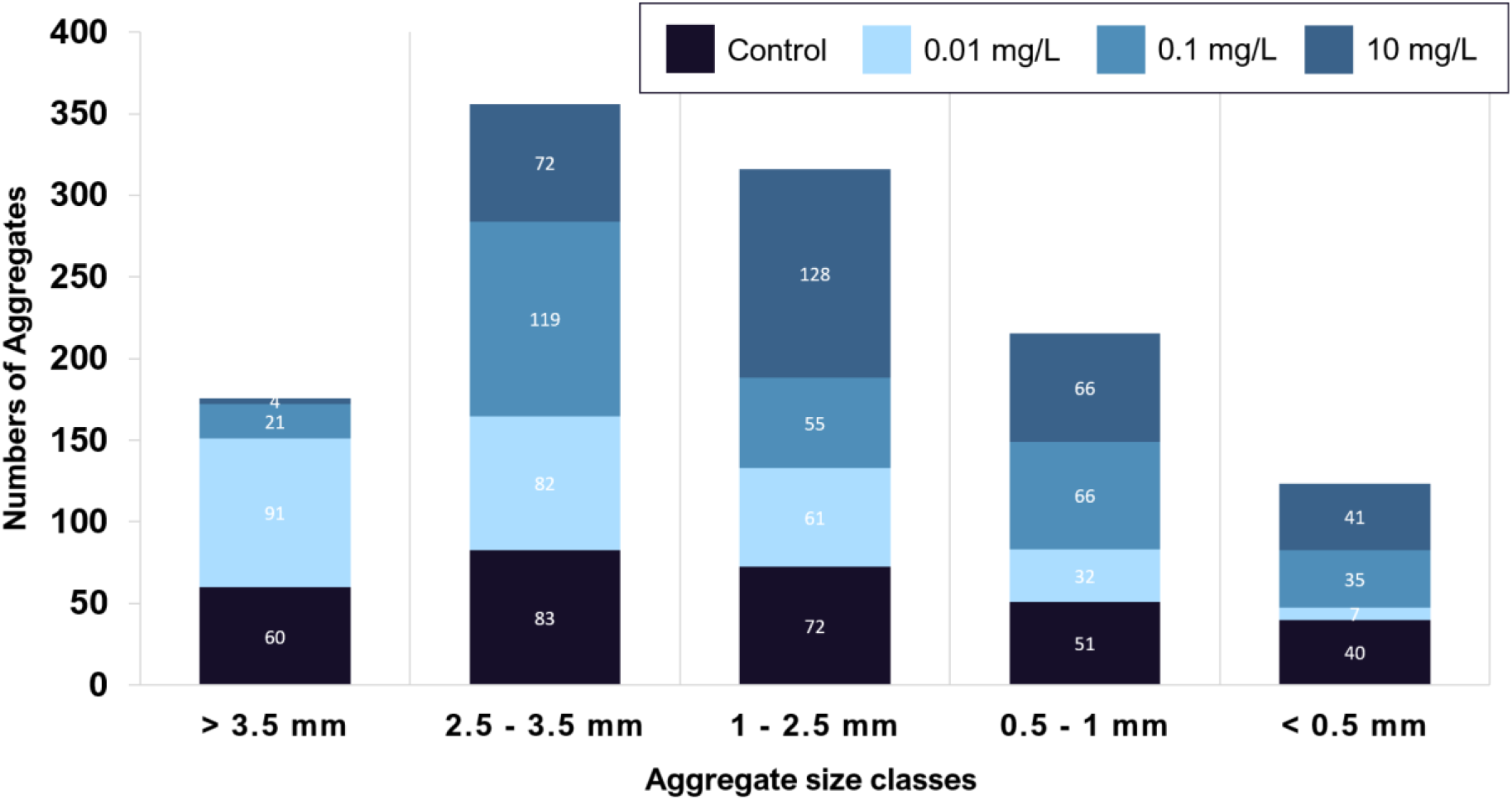
NanoPS exposure leads to change in aggregation. Aggregates sorted by size class show a significant change in distribution pattern under nanoPS exposure (Holm-Sidak, p = 0.05). The number of aggregates are reduced by 10 % in the 0.01 mg/L treatment (Holm-Sidak, p=0.003), but aggregation was enhanced overall in that treatment to have a higher percentage of huge aggregates than in the control treatment (Holm-Sidak, p = 0.001). In the higher plastic treatment at 10 mg/L this is reversed, leading to more aggregates overall, and more of those being of smaller sizes. No differences are observed when exposure length is compared.

Due to nanoPS exposure, aggregation and sinking velocities are impacted which in turn leads to change in sedimentation. As the majority of the host animals obtain their symbiotic dinoflagellates from the sand and water column [16], these changes in dinoflagellate sedimentation might lead to problems in acquisition of symbionts for the host animals. The lowest plastic treatment used, which is environmentally possible, already induces changes to the sedimentation. This lowest treatment led to bigger aggregates which at the same time sank faster, possibly removing the symbionts from the water column faster than required from the host animals and reducing chances of encountering symbionts.

**Figure 3.**
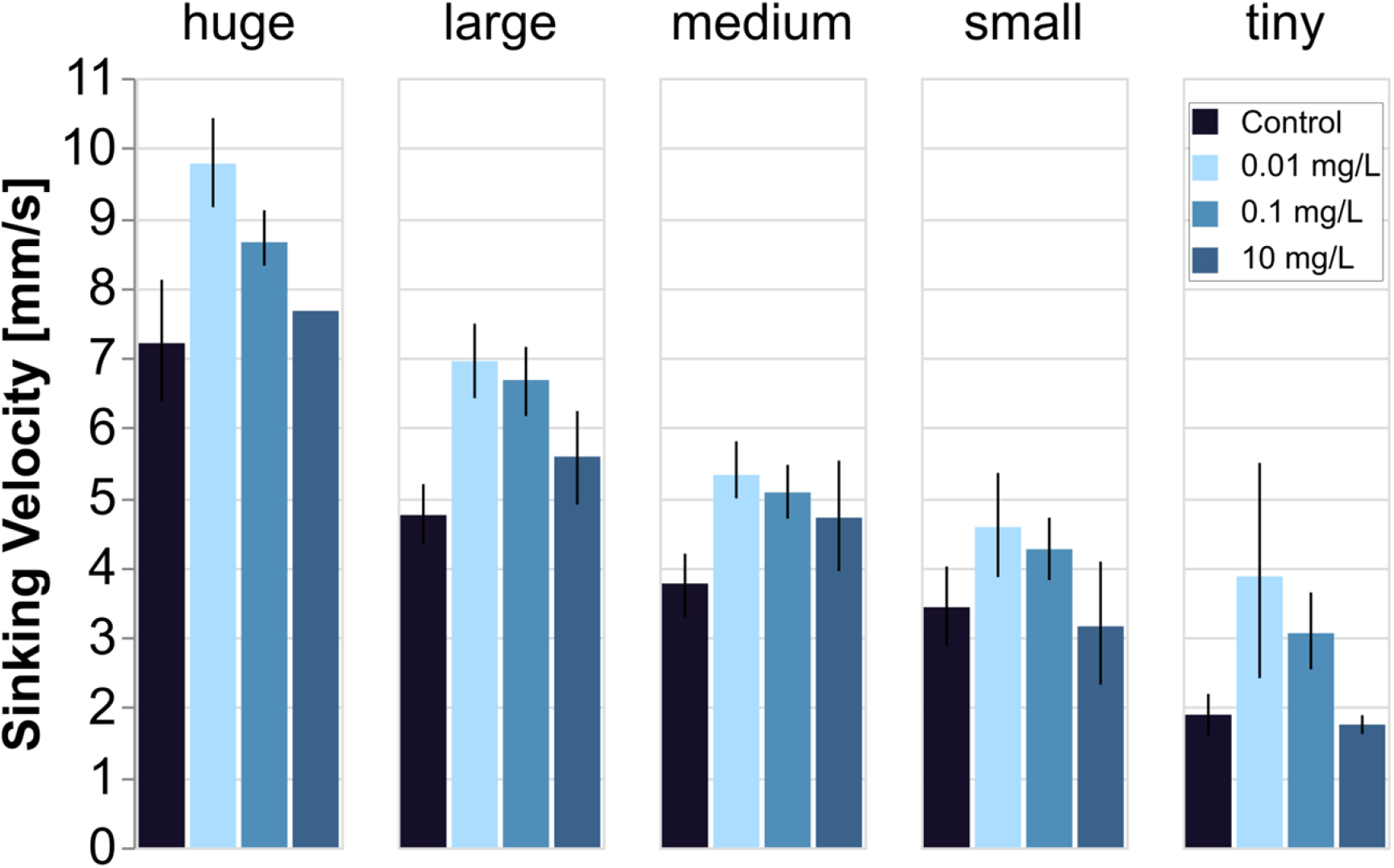
Sinking velocity change with nanoPS exposure. Sinking velocities decrease with aggregate size, from more than 7 mm/s (huge) to less than 2 mm/s (tiny). In all size classes, the control was similar in sinking velocity to the highest nanoPS treatment (10 mg/L). The low nanoPS treatment (0.01 mg/L) differed significantly from both controls (t-test, two-tailed p = 5.56 × 10^-4^) and the highest nanoPS treatment (t-test, two-tailed p = 9.03×10^-4^). This was also true for the intermediate nanoPS treatment (darker blue, 0.1 mg/L). Error bars are 95 % confidence intervals. Only one huge aggregate was measured in the highest nanoPS treatment. No differences in sinking velocity were observed in relation to exposure length.

Changes in aggregation and resulting sedimentation was observed under nanoPS exposure. It is interesting to see that the biggest changes in sinking velocity correspond to increases in aggregation and are observed in the lowest plastic treatment at 0.01 mg/L. On the other hand, the 10 mg/L treatment did not have any significant effect on the sinking rates but did affect sedimentation indirectly through changes in the aggregate size distribution (see Figure 2). These changes, both sinking velocities and aggregate sizes distribution, are most likely due to hetero-aggregation between algae and nanoPS. Under different treatments, the size distribution of aggregates was significantly different (see Figure 2). In combination, it is likely that the same effect that led to that difference in aggregation is also responsible for the difference in sinking velocities. Changes in EPS production and stickiness will lead to different cell packaging within the aggregates, possibly creating tighter packed aggregates in the lowest and intermediate treatment. This effect might be counteracted under the highest nanoPS exposure, by the sheer volume of EPS, which is lighter than seawater (Mari2017). The nanoplastic itself trapped in these could also add to the sinking velocity returning back to control levels in the high plastic treatments. As these symbionts are paired with the mobile larvae of the host animals, a higher sinking velocity would remove the potential symbiont from the pelagic area and reduce the chance of a match.

### 3.3. NanoPS effects on gene expression

Analysis of differential gene expression showed that in *Symbiodinium*, 14 genes were upregulated after nanoPS_42_ exposure, and 34 were downregulated relative to controls (Figure 2*a*). In *Cladocopium*, 75 genes were upregulated, and 169 genes were downregulated (Figure 2*b*). *Cladocopium* seems more sensitive to nanoPS_42_ exposure, as overall more genes responded than in *Symbiodinium*. Since Pfam analysis had more annotations than BLAST2GO in DEGs of *Cladocopium*, we list the major domains encoded by the DEGs of *Cladocopium*. (Supplementary Tables S3-S6).

The largest group of upregulated genes was a subfamily of dynein-related proteins having an AAA_5 domain (Table 1). Dynein is a microtubule-associated motor protein. Ten genes for dynein-related proteins with AAA and/or DHC (Dynein heavy chain) were upregulated in *Cladocopium* by nanoPS_42_ (Table 1, Supplementary Table S4). It has been shown that microplastic exposure induces production of reactive oxygen species (ROS) in microalgae [13,28] and dynein upregulation, therefore, it might be needed to balance cytoskeletal dynamics as microtubule polymerization is impaired by oxidative stress [33]. Interestingly, dynein light chain genes were also shown to be upregulated in gill cells of zebra mussels exposed to polystyrene microplastic [34].

**Table 1.** Domains encoded by more than three up-regulated genes in *Cladocopium* sp.

| Domain name | Summary from Pfam database | Gene number |
| --- | --- | --- |
| AAA_5 | AAA domain (dynein-related subfamily) | 6 |
| DHC_N2 | Dynein heavy chain, N-terminal region 2 | 5 |
| AAA | ATPase family associated with various cellular activities | 4 |
| AAA_6 | Hydrolytic ATP binding site of dynein motor region | 4 |
| TIG | IPT/TIG domain | 4 |

Four upregulated genes in *Cladocopium* (Table 1) encoded proteins with TIG domains that have an immunoglobulin-like fold and are found in cell surface receptors that control cell dissociation [35,36]. This might contribute to adhesion between neighboring cells and to the extracellular matrix composition, and explain some of the changes observed in cell aggregations.

There were more downregulated genes than upregulated genes in both *Symbiodinium* and *Cladocopium* (Figure 2). PPR (pentatricopeptide repeat) protein (Table 2) is involved in RNA editing [36] and extensive RNA editing has been reported in organelles of Symbiodiniaceae [37,38]. Five genes for photosynthesis were downregulated (Figure 2). These changes may explain observed reductions in photosystem efficiency in *C. goreaui* [13].

**Figure 4.**
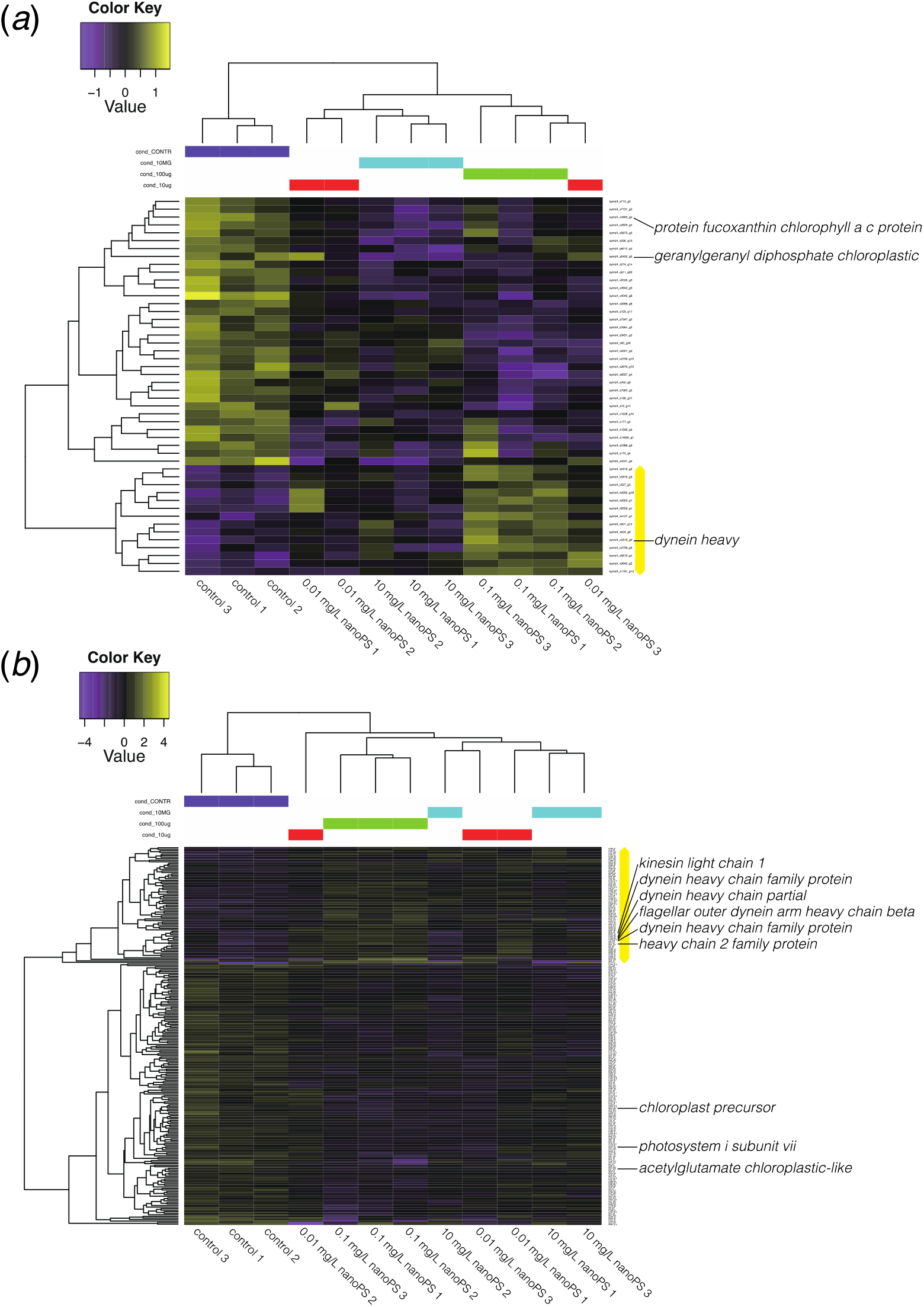
Heatmap and clustering of differentially expressed genes (2-fold changes, P<0.001) between dinoflagellates exposed to nanoplastics and controls. (*a*) DEGs in *Symbiodinium tridacnidorum*. (*b*) DEGs in *Cladocopium* sp. Values indicate the relative gene expression level, with purple and yellow showing downregulation and upregulation, respectively. The yellow bar shows a cluster of upregulated genes. Annotations by Blast2GO show the presence of microtubule- or photosynthesis-related genes among DEGs.

**Table 2.** Domains encoded by more than three down-regulated genes in *Cladocopium* sp.

| Domain name | Summary from Pfam database | Gene number |
| --- | --- | --- |
| Ank | Ankyrin repeat | 10 |
| Ank_2 | Ankyrin repeats (3 copies) | 10 |
| Ank_3 | Ankyrin repeat | 10 |
| Ank_4 | Ankyrin repeats (many copies) | 10 |
| Ank_5 | Ankyrin repeats (many copies) | 10 |
| PPR_2 | PPR repeat family | 6 |
| RCC1_2 | Regulator of chromosome condensation (RCC1) repeat | 6 |
| ANAPC3 (Apc3) | Anaphase-promoting complex, cyclosome, subunit 3 | 5 |
| Pkinase | Protein kinase domain | 5 |
| PPR | PPR repeat | 5 |
| PPR_3 | Pentatricopeptide repeat domain | 5 |
| Abhydrolase_5 | Alpha/beta hydrolase family | 4 |
| Abhydrolase_6 | Alpha/beta hydrolase family | 4 |
| Lipase_3 | Lipase (class 3) | 4 |
| PPR_1 | PPR repeat | 4 |
| TPR_14 | Tetratricopeptide repeat | 4 |
| YukD | WXG100 protein secretion system (Wss), protein YukD | 4 |

Other downregulated gene groups were related to intracellular degradation processes, including hydrolase and lipase, and to subunit 3 of the anaphase-promoting complex/cyclosome [40]. The downregulated gene (s3282_g2) with abhydrolase and chlorophyllase domains is likely related to chlorophyll degradation [41]. The gene, s576_g21, for cell division control (CDC) protein 2 is downregulated in *Cladocopium*. Downregulation of six genes with RCC1 (regulator of chromosome condensation) and three genes with CDC domains suggest some effect on cell division. Thus, several negative consequences of nanoPS_42_ exposure are suggested by DEGs (summarized in Supplementary Figure S4).

## 4. Conclusions

Previous studies have shown that nanoplastic has adverse effects on different algae groups [27,29,30,42,43], and a recent study shows that microplastic has similarly negative effects on an endosymbiotic dinoflagellate *Cladocopium goreaui* [13]. No previous studies have been conducted on nanoPS_42_ effects on Symbiodiniaceae. We found significant changes in aggregation and aggregate sinking velocity of *Symbiodinium tridacnidorum* and *Cladocopium* sp., coupled with variations in gene expression patterns after exposure to nanoPS_42_. This suggests that nanoPS_42_ in coral reef ecosystems has the potential to influence the acquisition of symbionts by mollusks and corals, likely damaging these symbiotic relationships. Since both are major architects of reef structure, nanoPS_42_ pollution has the potential to lead to structural changes in reef ecosystem dynamics.

**Figure 5.**
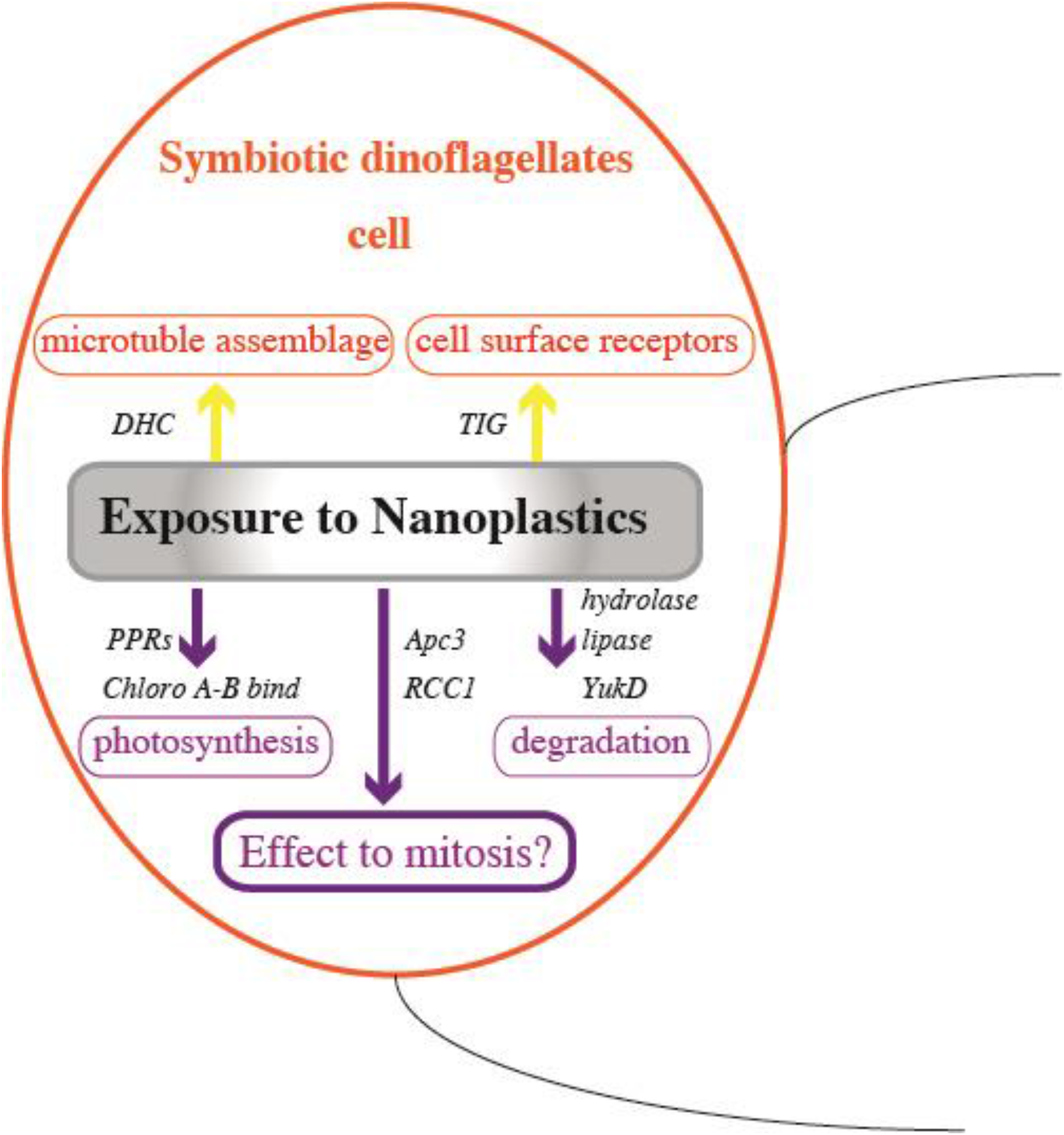
Exposure to nanoPS_42_ changes gene expression levels in symbiotic dinoflagellates. Yellow and purple arrows show up-regulation and down-regulation of gene expression, respectively.

## Supplementary Materials

The following are available online at www.mdpi.com/xxx/s1; Figure S1: Cell abundance in treatment tanks, control tanks, and outside controls; Figure S2: NanoPS exposure changes aggregation behaviour, reduces cell numbers, and alters size class distributions; Table S1: Relationship between nanoPS_42_ concentration and particles per Tank; Table S2: Sampling days of each tank; Table S3: Genes that responded to nanoplastic exposure in *Symbiodinium tridacnidorum;* Table S4: Genes that responded to nanoplastic exposure in *Cladocopium* sp.

## Author Contributions

CR designed the study and performed the experiment. ES carried out RNA analyses. KK and CR performed RNA-seq mapping and cluster analyses. All authors wrote the manuscript. All authors have read and agreed to the published version of the manuscript.

## Acknowledgments

We are grateful to the DNA sequencing section of OIST for RNA preparation and sequencing and to the OIST imaging section for 3D image support. We thank members of the MGU, especially Ms. Haruhi Narisoko for cell culturing and Ms. Kanako Hisata for IT support. We are grateful for the help and support provided by Dr. K. Deasy from the Engineering Support Section of Research Support Division at OIST. We gratefully acknowledge Profs. Síle Nic Chormaic and Noriyuki Satoh for their continuing personal and material support and kind encouragement throughout the project.

## Availability of Data

Data are available in the electronic supplementary material. Raw sequence data are available from PRJNA627564 in NCBI database. *Symbiodinium* (currently the family Symbiodiniaceae) clade A3 and C genomes: clade A3 (https://marinegenomics.oist.jp/symb/viewer/info?project_id=37) clade C (https://marinegenomics.oist.jp/symb/viewer/info?project_id=40) Transcript models of *Symbiodinium* clades A3 and C: https://marinegenomics.oist.jp/symb/download/syma_transcriptome_37.fasta.gz https://marinegenomics.oist.jp/symb/download/symC_aug_40.fa.gz

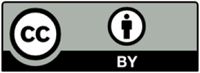 © 2020 by the authors. Submitted for possible open access publication under the terms and conditions of the Creative Commons Attribution (CC BY) license (http://creativecommons.org/licenses/by/4.0/).

